# The (implicit) force is with you: an application of the implicit function theorem for automatic protein force fields reparametrization

**DOI:** 10.64898/2026.10.11.758296

**Authors:** Satya Tamby, Ardite Bujupaj, Gabriele Orlando

## Abstract

Molecular dynamics simulations are powerful tools for studying protein behavior, yet their accuracy remains limited by the approximations of molecular force fields and the difficulty of determining their parameters. Moreover, existing force fields are largely optimized for globular proteins and typically perform poorly on structurally distinct systems. Developing a universal force field that accurately describes all existing protein structures may be beyond reach with current physical models. Instead, we propose shifting generality from the force field itself to the method used to assign its parameters, enabling systematic reparametrization for restricted, structurally homogeneous classes of proteins. We formulate this task as a bilevel optimization problem and solve it using the implicit function theorem. We demonstrate this approach by reparametrizing AMBER for amyloid fibrils, optimizing its parameters to improve agreement with experimentally determined structures and enhance fibril stability during molecular dynamics simulations.

## INTRODUCTION

Atomistic molecular dynamics simulations (MD) are among the most widely used computational tools in modern science. For decades they have been applied across chemistry, biology, and materials science, from proteins and nucleic acids to membranes and polymers. At the base of their functioning there is a set of energy functions, commonly called force fields. A force field provides the mathematical description of how atoms in a molecular system interact. Given a set of atomic coordinates, it evaluates their interactions in order to compute the potential energy of the system along with of the forces acting on each atom. By iteratively updating atomic positions according to these forces over small time steps, the system evolves in time, thereby enabling molecular dynamics simulations that sample conformational changes and thermally accessible states of the molecule [1].

A particularly important class of force fields is the one developed for proteins, where molecular dynamics simulations are suited to capture the flexibility and the conformational changes characteristic of many protein systems. These protein force fields assign a set of parameters to the atoms of amino acids, providing the numerical values required by the underlying equations that describe the molecular interactions. These values determine how the general mathematical form of the equations is translated into the specific energetic and force behavior of a given molecular system. Among the most widely used are the AMBER family of force fields, which provide standardized parameter sets for simulating proteins [2, 3].

At a more fundamental level, however, all atomistic force fields rely on an unavoidable simplification of interatomic interactions. The true potential energy surface of molecular systems is inherently many-body, involving a countless number of variables. Force fields usually replace this complexity with pairwise additive terms and fixed functional forms, which stay fixed regardless of the chemical environment or the context in which two interacting atoms are found. On top of this, many of the parameters and of the coefficients of these terms cannot be measured directly and are, instead, inferred from simplified systems that are easier to study experimentally or computationally. For example, the strength of attractive interactions between atoms is often adjusted to reproduce properties of small molecules, such as their solubility, or tendency to remain in a specific liquid phase [4, 5]. Similarly, atomic charge distributions are commonly derived from quantum mechanical calculations on isolated molecules [6]. These parameters are subsequently transferred to much larger and more complex systems, under the assumption that the same effective interactions remain valid across different chemical environments. As a result, force field parameters often reflect behavior observed in simplified reference systems rather than the full complexity of the biological systems to which they are ultimately applied [7]. This reductionist approach is essential for computational tractability, but it also means that the resulting models necessarily rely on substantial approximations.

These limitations become increasingly pronounced as the complexity of the system increases. In proteins, achieving agreement with experimental observations often requires successive empirical corrections, parameter adjustments, and specialized refinements to compensate for deficiencies in the underlying force field and maintain physically realistic behavior [8], particularly for proteins that fall outside the classical globular protein paradigm.

Amyloid fibrils are one of them: they are a protein-based superstructure which are characterized by an exceptional thermodynamic stability, often described as effectively irreversible under physiological conditions, and they can resist disassembly even under extreme perturbations such as changes in temperature or pH [9]. Despite this experimental robustness, conventional molecular dynamics simulations based on standard force fields frequently fail to reproduce this behavior. Instead, they often predict that experimentally resolved fibrillar structures are unstable and that they spontaneously dissociate into monomers or disordered aggregates [10, 9]. This indicates a systematic mismatch between the modeled energetic landscape and the experimentally observed stability of these systems.

Efforts to improve force fields have largely focused on patching individual equations or introducing targeted corrections when discrepancies with experiments are observed [11]. In this sense, force field development has often relied on iterative tuning guided by expert knowledge, rather than on a global and data-driven optimization of the model as a whole. Although these adjustments often improve agreement for specific benchmarks, they do not address the fact that inaccurate parameters remain embedded within a simplified model.

Accepting that force fields are necessarily simplified descriptions of reality suggests a different strategy: rather than seeking a universal parameter set capable of accurately describing every possible molecular system, it may be more realistic to develop models tailored to well-defined classes of molecules that share similar structural and physicochemical properties. For proteins, for example, one could focus on homogeneous families with related folds and interaction patterns, and infer the parameters that best describe the experimental behavior of that specific ensemble. Such an approach should ideally be generalizable and applicable to any protein subgroup in an agnostic, data-driven manner, without requiring manual parameter adjustments. This strategy should leverage an experimental data source which is extensively available for most protein families, such as high-resolution protein structures.

In this article, we explore an approach which has the objective to identify the force field parameters for which the experimentally observed structures, which we consider statistical approximations of lowest-energy conformations, are placed in an energetic minimum of the energy landscape defined by the force field.

This naturally defines two nested optimization problems: in the inner problem, the atoms are moved to find the conformation at the energetic minimum under the current parameters. The outer problem adjusts the parameters so that this conformation coincides with the experimental one. Iterating the two problems progressively reshapes the energy landscape until its minima coincide as closely as possible with the experimentally observed structures (Figure 1). This class of problems, in which the variables of one optimization are determined by the solution of another, is known as bilevel optimization and has been extensively studied in applied mathematics [12].

**Figure 1:**
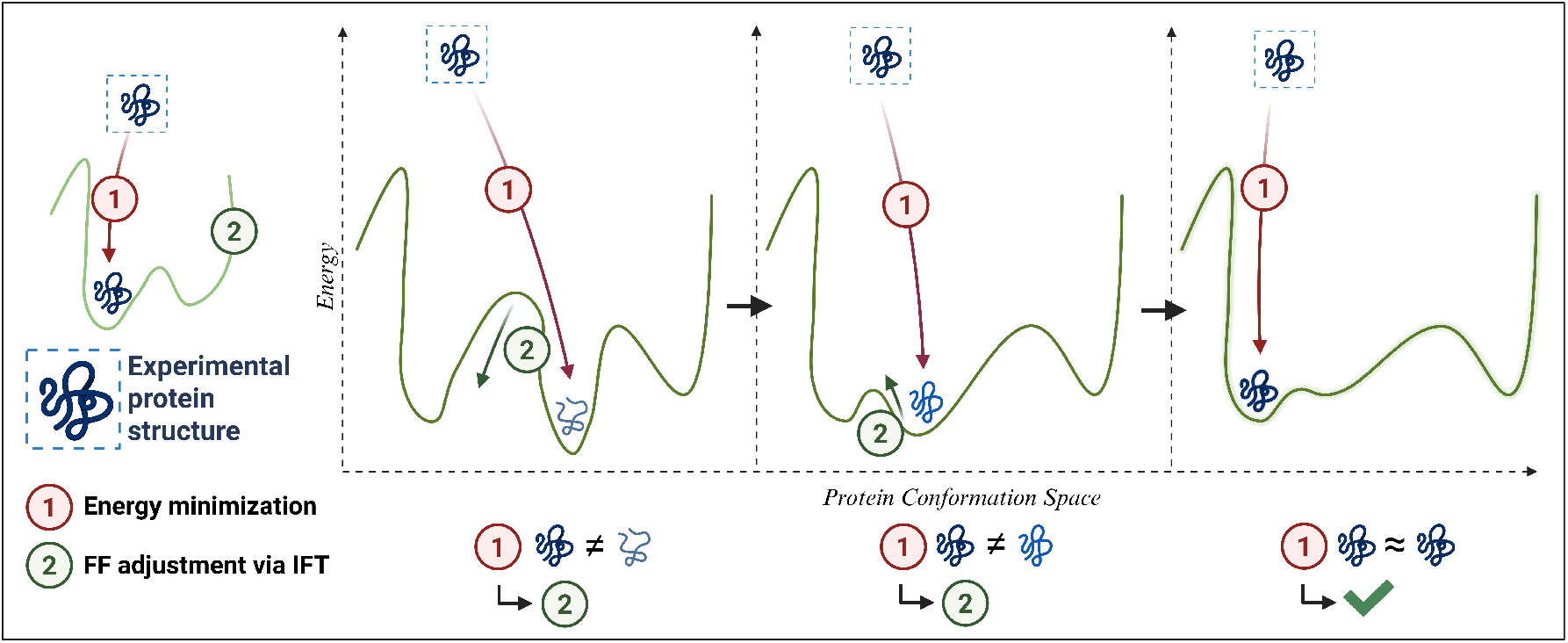
Bilevel optimization scheme for force field reparametrization. Starting from an initial set of structures and the initial force field F (**θ**_initial_), the parameters **θ** of the force field are optimized through two nested loops. In the inner loop (1), the structure is relaxed by energy minimization under the current parameters: the energy and forces are evaluated, and the coordinates are updated until a stationary point is reached, providing the relaxed conformation. In the outer loop (2), this relaxed conformation is compared to the experimental one through the loss. The gradient of the loss with respect to the parameters is obtained via the implicit function theorem and is used to update **θ**. Iterating the two loops drives the relaxed structure toward the experimental one and returns the reparametrized force field F (**θ**_final_). FF, Force Field; IFT, Implicid Function Theorem

The key methodological advantage of this formulation is the possibility of using the implicit function theorem (IFT) [13, 14]. This theorem allows the gradient of the loss, which defines how far the minimum of the force field is from the experimentally determined structure, with respect to the force field parameters to be computed directly from the converged minimum, without differentiating through the individual steps of the minimization process. The force field parameters can therefore be optimized with standard gradient-based methods, rather than with gradient-free searches whose cost grows rapidly with the number of parameters. Because only the converged minimum is needed, the cost of computing the gradient is independent of the number of minimization steps, which is critical when each energy minimization can require hundreds of thousands of iterations, as is typical for protein systems. Moreover, the energy minimizer can be treated as a black box: only the final minimized structure and the derivatives of the force field at that point are required. This makes it possible to reparametrize the force field while relying on established, highly optimized simulation engines such as OpenMM [15] to carry out the minimization, instead of reimplementing an *ad hoc*, fully differentiable simulator.

To prove the effectiveness of this approach, we use it to adjust the parameters of the AMBER force field, focusing on a structurally distinct and challenging class of biomolecular assemblies: amyloid fibrils. We found that a mild reoptimization of parameters, limited to deviations of approximately 30% from their original values, is sufficient to correct major macroscopic simulation artifacts observed with standard parameter sets. These improvements are obtained through a fully data-driven and agnostic reparametrization procedure, without explicitly encoding or targeting any of the problems affecting amyloid structures, but instead allowing it to emerge naturally from the optimization against experimental structural constraints. Therefore, the same class-specific reparametrization strategy can be applied to any molecular family for which sufficiently large sets of high-resolution structures are available.

## RESULTS

### Amyloid structures reveal widespread instability under standard force fields

Amyloid proteins represent a class of structures that differs substantially from the globular proteins on which current force fields were largely optimized. Amyloids are poorly soluble, highly repetitive supramolecular assemblies that have only recently become accessible at atomic resolution through cryoEM [16]. Today, a few hundred amyloid structures are available in the PDB, but these data are highly heterogeneous. Structures often contain different numbers of fibril layers, despite amyloid fibrils being, in principle, infinitely extendable, and many contain missing atoms or regions with poorly resolved atomic coordinates. To obtain a homogeneous dataset, we used Amyloid Explorer [17, 18], a manually curated collection of amyloid structures from which we collected 121 amyloid structures (see Methods for details).

We first evaluated the behavior of standard molecular mechanics force fields on this dataset. Each structure was subjected to energy minimization using the AMBER force field parameterization. Our objective was to quantify how closely the minimum energy conformation predicted by the force field matched the experimentally determined structure. The resulting median root mean squared deviation (RMSD) between the minimized structures and the experimental structure is 4.11 Å (Figure 2D). Structural drift was often substantial, with 47 cases showing an RMSD greater than 5 Å from the experimental structure.

**Figure 2:**
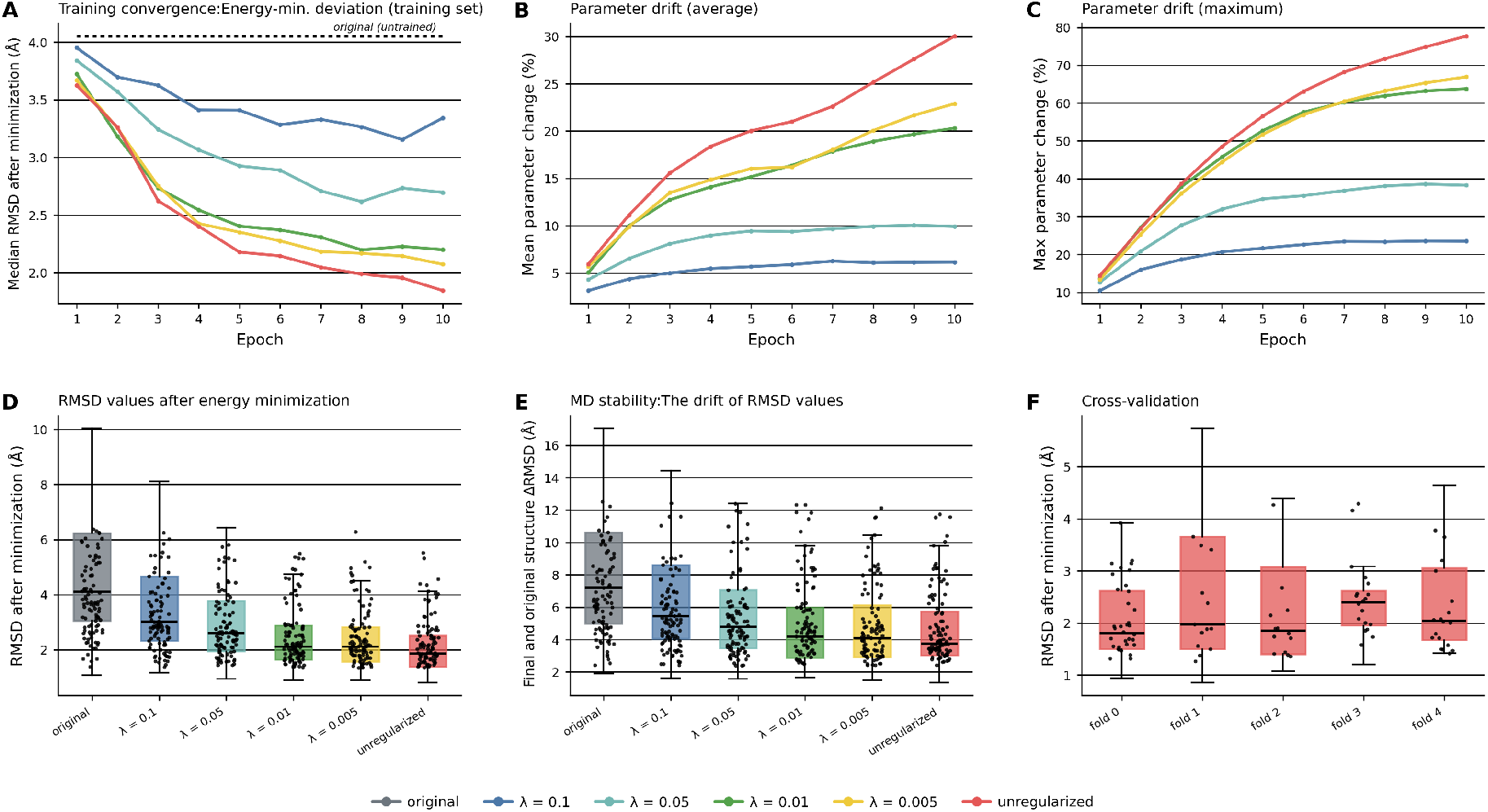
**A)** Convergence of the IFT-based reparametrization using different values of λ. The plot shows the evolution of the RMSD between the energy-minimized structures and their corresponding experimental conformations during training, using different regularization strengths. As expected, increasing the regularization coefficient λ progressively constrains parameter updates, making convergence toward lower RMSD values more difficult. **B)** Average deviation from the original force field at different regularization levels. The figure shows the evolution of the average parameter change during training for different values of λ, which controls the strength of the regularization. As expected, increasing λ progressively constrains the parameter updates, reducing the overall deviation from the original force field. The average parameter change decreases from approximately 30% in the unconstrained optimization to approximately 6% when using λ = 0.1. **C)** Maximum deviation from the original force field at different regularization levels. The figure shows the evolution of the maximum individual parameter change during training for different values of λ, which controls the strength of the regularization. Increasing λ progressively limits large parameter updates, reducing the maximum deviation from the original force field. The maximum parameter change decreases from above 70% in the unconstrained optimization to approximately 24% when using λ = 0.1. **D)** RMSD after energy minimization across the amyloid dataset. Distribution of the RMSD between the energy-minimized and experimental structures over the 121 amyloid structures, for the original AMBER force field and the five reparametrized variants. **E)** MD stability. Distribution of the structural drift (RMSD between the final frame of the 0.5 ns MD simulation and the experimental structure) over the amyloid dataset, for each force field variant. **F)** Cross-validation. RMSD after energy minimization for each of the five cross-validation folds (models trained without regularization λ = 0) on four fifths of the dataset and evaluated on the remaining fifth. All RMSD values are reported in Å.

Energy minimization, however, represents only a limited application of these force fields, which are primarily designed for molecular dynamics simulations. We therefore performed 0.5 ns molecular dynamics simulations for every amyloid structure in the dataset. The median RMSD of the final simulated conformations with respect to the experimental structures is 7.20 Å. These results indicate that amyloid fibrils, which are experimentally known to be exceptionally stable supramolecular assemblies, are systematically predicted to be unstable by current force fields. In a non-negligible fraction of cases (37 out of 121 cases), fibrils spontaneously disassemble within only a few tens of picoseconds (final state RMSD with respect to original conformation greater than 10 Å). These predictions are clearly inconsistent with experimental observations and suggest that current force fields might have issues in defining the behavior of this important class of proteins. We also verified that this behaviour is not an artefact of the short simulation time: extending the simulations tenfold, to 5 ns, left the median drift of the original force field essentially unchanged (7.80 Å at 5 ns versus 7.20 Å at 0.5 ns).

### Optimization of the force field parameters using the implicit function theorem

Force field parameter optimization can be formulated as a two-step process involving an inner optimization and an outer optimization:

The inner optimization corresponds to the standard energy minimization procedure and its objective is to find the coordinates that minimize the potential energy. Assuming that the force field is differentiable, this minimum can be found using gradient-based optimization. This is the approach implemented, for example, in OpenMM for energy minimization.

The outer optimization adjusts the force field parameters. The objective is to find the parameters that minimize the root-mean-square deviation between the relaxed structure returned by the inner problem and the experimental structure.

Every update of the parameters therefore requires solving a complete energy minimization problem to obtain the corresponding relaxed structure. This creates a nested optimization procedure.

At first sight, this dependency appears to prevent the use of conventional gradient based optimization, since the optimization variables of the outer problem depend on the solution of another optimization problem. However, by applying the implicit function theorem, it can be shown that, under appropriate conditions, the gradient of the outer objective with respect to the force field parameters does not depend on the optimization trajectory followed by the inner minimization. Instead, it depends only on the final relaxed conformation. The key requirement is that the relaxed structure corresponds to a stationary point of the energy, that is, ∇_*X*_ *E* ≈ 0.

The complete derivation of the optimization equations is presented in the Methods section. This formulation has two important consequences. First, the parameter optimization becomes independent of the specific algorithm used to perform the energy minimization. Second, the minimization procedure can be treated as a black box, provided that it returns a relaxed structure for a given set of force field parameters. This allows the direct integration of highly optimized external software, such as OpenMM, without requiring differentiation through the minimization algorithm itself.

We applied the proposed bilevel optimization framework to the amyloid dataset with the objective of reparametrizing the AMBER force field to improve the agreement between energy minimized structures and their experimentally determined conformations. The trainable parameters were initialized with the original AMBER99SB values and iteratively optimized.

At each training iteration and for each amyloid structure, an energy minimization step was first performed using OpenMM with the current set of parameters (inner optimization). Once the relaxed structure corresponding to the minimum of the potential energy was obtained, its RMSD with respect to the experimental structure was computed. This RMSD served as the outer objective. The gradient of the RMSD with respect to the force field parameters was then computed using the implicit function theorem (see Methods). The resulting gradient was used by a standard Adam optimizer to update the force field parameters. The process was repeated for every amyloid in the dataset, and then iterated 10 times.

The median RMSD over the training iterations decreases uniformly with the optimization epochs, showing that the IFT-based training consistently produces conformations that are substantially closer to the experimentally determined conformations than the original force field.

To evaluate whether the optimized parameters also improve molecular dynamics simulations, we performed MD simulations using the reparametrized force field. The simulation protocol was identical to that used with the original AMBER force field, with the only difference being the optimized values of the parameters. The results show that the drift from the experimental structure passes from 7.20 Å to 3.76 Å (Figure 2E). As for the original force field, this value remained essentially unchanged when the simulations were extended tenfold to 5 ns (4.33 Å). Moreover, simulations performed with the original force field frequently resulted in complete amyloid fibril disassembly. In contrast, the reparametrized force field substantially improved fibril stability, with only 9 cases showing a drift greater than 10 Å over the same simulation time. Figure 3 shows a representative comparison between molecular dynamics simulations performed with the original AMBER force field and the reparametrized version for the tau amyloid structure (PDB ID: 8G54). The original AMBER force field predicts rapid dissociation of the amyloid assembly, whereas the reparametrized force field preserves the integrity of the fibril throughout the simulation. This example qualitatively illustrates the improved stability achieved by the optimized force field and is representative of the behavior observed across the broader amyloid dataset.

**Figure 3:**
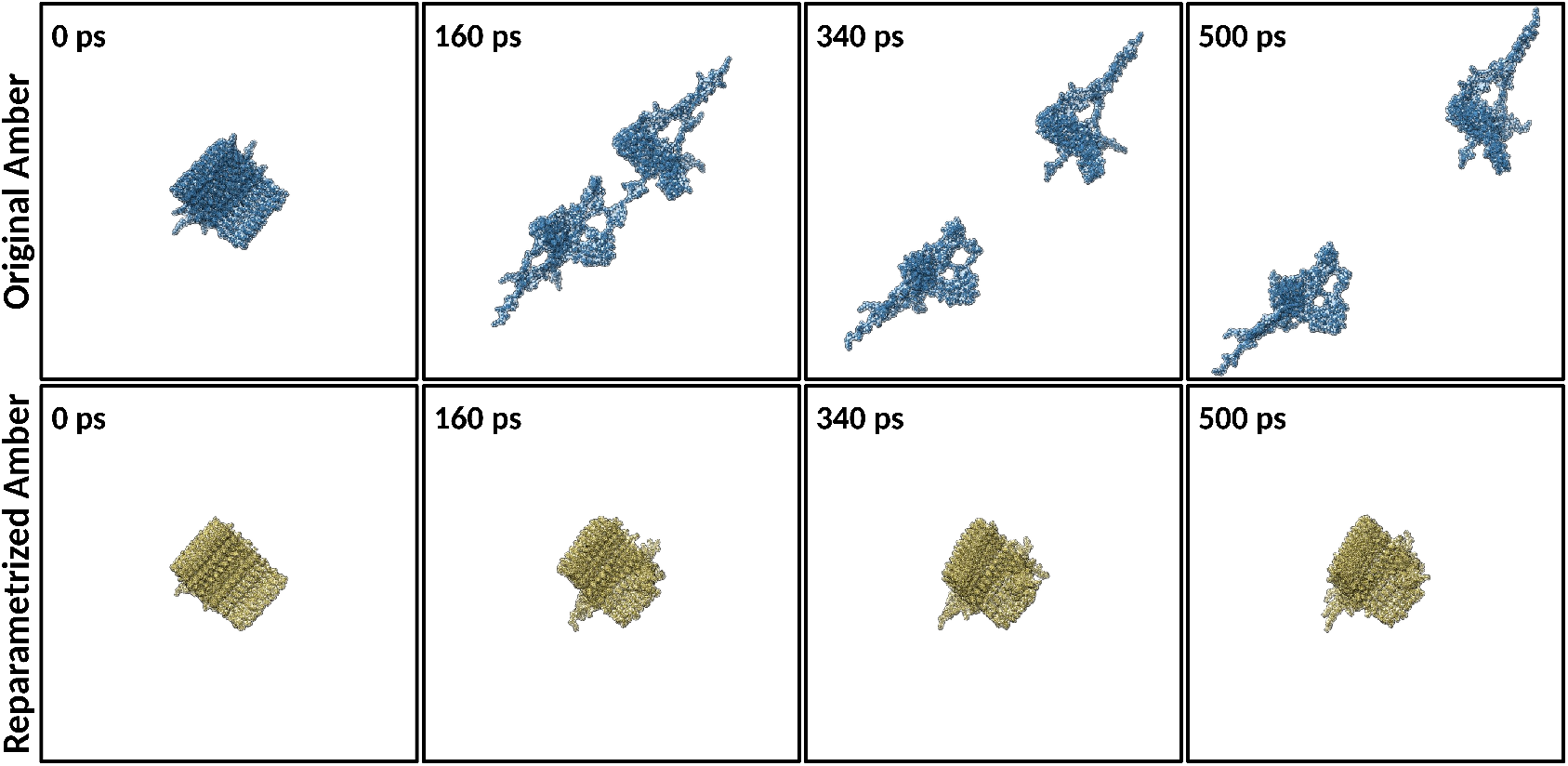
Comparison of MD simulations using the original and reparametrized AMBER force fields. The figure shows representative snapshots from molecular dynamics simulations of the tau amyloid structure (PDB ID: 8G54) performed with the original AMBER force field (top row) and the reparametrized force field (bottom row). While the original AMBER parameterization leads to rapid disassembly of the amyloid assembly, the reparametrized force field preserves the structural integrity of the fibril throughout the simulation.

To verify that this improvement reflects a genuine, generalizable refinement of the force field rather than overfitting to the specific structures used for training, we performed a five-fold cross-validation: the force field was reparametrized on four fifths of the amyloid dataset and evaluated on the remaining fifth (Figure 2F). To ensure that this generalization was not biased by redundant structures (sequence polymorphs, or different resolved ranges of the same protein) between the training and test sets, the folds were defined by grouping: structures were clustered by sequence (two structures placed in the same cluster whenever one monomer sequence is a substring of the other), and whole clusters were assigned to a single fold, so that no protein, and no other member of its family, ever appeared in both the training and the test set. Across the five folds, the median RMSD on the testing structures is 1.97 Å, essentially identical to the value obtained when training on the full dataset (1.86 Å) and well below the original AMBER99SB baseline (4.11 Å).

These results suggest that bilevel optimization can effectively refine a molecular mechanics force field for a specific class of proteins. Importantly, the optimization is performed solely on energy minimized structures, without directly optimizing molecular dynamics trajectories. Nevertheless, the learned parameters generalize to MD simulations, producing trajectories that are substantially more consistent with the experimentally observed amyloid structures.

### Minimum changes for maximum outcome

While the optimization procedure successfully improves the agreement between the relaxed and experimentally determined amyloid structures, we next investigated how this improvement was achieved at the level of the force field parameters. Starting from the original AMBER99SB parameterization, the optimization leads to an average parameter variation of 30.0% after 10 training epochs. However, this global change hides larger modifications occurring for individual parameters, with the maximum deviation reaching 77.7%. An excessively aggressive modification of the force field may not always be desirable: large deviations from the original parameterization could break the physical laws encoded in the force field, resulting in physical inconsistencies. Therefore, an ideal optimization should not only improve structural agreement but should also preserve, as much as possible, the original parameterization.

To achieve this balance, we introduced a regularization term that penalizes large deviations of the optimized parameters from their initial AMBER99SB values. This approach is commonly used in machine learning, particularly in neural network training, where it is referred to as weight decay and is used to prevent individual parameters from reaching extreme values. In this context, we applied an L2 regularization, which penalizes parameter changes proportionally to their squared magnitude. As a consequence, small parameter variations are only weakly penalized, while increasingly larger deviations are progressively discouraged. The strength of this regularization is controlled by a coefficient, referred to as *λ*.

The RMSD evolution obtained with different values of *λ* is reported in Figure 2A, while the corresponding changes in the average parameter variation and in the maximum individual parameter deviation are shown in Figures 2B and 2C, respectively.

As expected, increasing the regularization strength progressively limits the extent to which the force field parameters can deviate from their original AMBER99SB values. This effect is observed both in the average parameter variation and in the maximum change of individual parameters, suggesting that *λ* effectively controls the degree of freedom of the optimization process. However, constraining the parameter updates too strongly comes at the cost of reduced optimization efficiency. In particular, high regularization values (*λ* = 0.05 and *λ* = 0.1) excessively restrict the parameter space explored by the optimizer, impairing convergence significantly.

In contrast, milder regularization provides a better balance between structural accuracy and parameter conservation. For smaller values of *λ*, the optimization converges to RMSD values between the relaxed and experimental structures of approximately 2.1 Å (Figure 2D), comparable to the unregularized optimization. At the same time, the required parameter modifications are substantially reduced. For example, with *λ* = 0.01, the overall parameter variation is limited to 20.3%, compared with 30.0% for the unregularized optimization, while maintaining only a marginal difference in agreement with experimental structures. The effect is even more pronounced when considering the largest individual parameter modification: the maximum change decreases from 77.7% in the unregularized case to 63.8% with *λ* = 0.01.

These results show that L2 regularization can be effectively applied to molecular force field refinement. By limiting excessive parameter modifications while preserving the ability to improve agreement with experimental structures, regularization provides a practical approach to obtaining optimized force fields that remain close to the original physical parameterization.

### Robustness of the reparametrized minima to structural perturbations

During training, each inner energy minimization was initialized from the corresponding experimental structure. This was a deliberate computational choice, as starting from random conformations would place the system farther from the target conformation and slow down training.

This choice, however, raises an important question: is the optimization improving the energy landscape, or is it simply producing a force field under which energy minimization causes minimal displacement from the starting structure? An effective reparametrization should not only place the energy minimum closer to the experimental conformation, but also retain the ability to relax perturbed structures toward its minimum.

To investigate this, we performed a test on 121 amyloid structures. We perturbed the atomic positions of each experimental structure by adding Gaussian noise, resulting in an initial displacement of approximately 4 Å RMSD. The same perturbed structure was used as the starting point for minimization with both force fields.

We then measured the total atomic displacement during minimization, defined as the RMSD between the perturbed starting structure and the resulting minimum. In other words, this test was meant to check if the optimization was actually actively moving atoms after the reparametrization. Both force fields produced substantial structural changes, with median displacements of approximately 6.7 Å for the original AMBER force field and 5.5 Å for the reparametrized force field. These results show that the reparametrized force field remains active and does not achieve improved agreement with the experimental structures simply by suppressing atomic movements during energy minimization.

### Reparametrised force fields preserve basic physics

As discussed above, developing a universal force field capable of accurately describing all classes of proteins and conformational states, including both globular and non-globular assemblies, remains an extremely challenging goal given our current understanding of protein physics. However, any meaningful force field refinement should preserve the fundamental physical principles required to maintain the stability of well-characterized protein folds. A reparametrization that improves the description of amyloid structures but causes small globular proteins to lose their native conformations would likely indicate that the optimization introduced unphysical distortions and might be the result of overfitting or other dataset-related biases.

To assess whether the optimized force fields preserve general protein stability, we evaluated their behavior on a dataset of 57 small globular proteins obtained from [19]. For each structure, we performed energy minimization using the original AMBER force field and the reparametrized versions obtained with different regularization strengths. We then compared the minimized structures to their experimentally determined conformations by evaluating whether any force field variant introduced unrealistic structural deviations, such as complete unfolding or loss of the native fold.

The results are reported in Supplementary Figure S1. All reparametrized force fields produced relaxation behavior comparable to the original AMBER parameterization, with no evidence of systematic destabilization of globular proteins. The RMSD distributions obtained after minimization were highly similar across all force field variants, indicating that the modifications introduced through the implicit function theorem-based optimization do not disrupt the fundamental physical properties required for maintaining stable protein folds. These results support the conclusion that the learned parameter changes are not simply a consequence of overfitting to amyloid structures, but instead represent targeted adaptations that improve amyloid description while preserving general protein stability. The same conclusion holds under molecular dynamics: applying the identical 0.5 ns MD protocol to the globular set, the reparametrized force fields produced structural drift comparable to the original AMBER parameterization (median drift in the range 5.6–7.3 Å across variants, versus 5.7 Å for the original), with no systematic destabilization.

### Reparametrization Targets Specific Interactions

We next analyzed which parameters are most responsible for these changes, and we assessed that the reparametrization primarily affects atomic partial charges. In the unregularized model, charge multipliers change by 57% on average. Moreover, the ten most strongly modified parameters are all associated with atomic charges (Supplementary Fig. S2). The largest changes occur for carboxylate oxygens (atom class O2, 73%) and hydrogens on carbon adjacent to positively charged groups (HP, 78%), followed by several further backbone and side-chain classes (H, HC, N, and S), which change by 59-64%.

This pattern is largely preserved across the different regularization we tested. Although the magnitude of the parameter changes decreases with increasing *λ*, the identity and relative ranking of the most strongly affected charge classes remain similar. The same pattern is also observed across different training subsets. In the five-fold cross-validation the per-class charge changes are highly consistent across folds.

The consistency of these changes across both regularization strengths and training subsets suggests that the reparametrization is driven by a reproducible signal in the data rather than by random variation.

#### Charge redistribution versus atom-class abundance

Our analysis revealed that the parameters undergoing the largest modifications during reparametrization are primarily associated with atomic charges. To determine whether this charge redistribution reflects a chemically meaningful optimization rather than a simple consequence of atom-class abundance in the dataset, we tested whether the frequency of each atom class influenced the magnitude of the corresponding charge changes.

Of the 44 atom classes defined by AMBER99SB, 29 are actually present in this protein dataset (the remainder correspond to nucleic-acid and monoatomic-ion classes and never occur), and the abundance analysis was restricted to these 29 classes. For each class, we compared its total abundance to the magnitude of its charge modification using the Spearman coefficient, computed separately for each of the five trained variants.

No significant relationship was observed between atom-class frequency and charge variation (Supplementary Fig. S3): the correlation coefficients remained close to zero, indicating that the optimization is not simply driven by the number of occurrences of a given atom type in the dataset.

#### Sequence specificity: shuffled-sequence decoys

As a further check, we investigated whether the reparametrized force fields retain sequence specificity, rather than nonspecifically driving every structure toward its experimental conformation.

To test whether this bias existed, we wanted to test whether the method was behaving differently when fed with sequence-shuffled decoy instead of experimental amyloid structures. Within each structure, the amino-acid identities were randomly permuted while the experimental backbone geometry was left unchanged. Chains sharing an identical sequence received the same permutation so that the fibril symmetry was preserved. The backbone atoms (N, C*α*, C, O) were never moved, and side chains were rebuilt with PDBFixer to match the new residue identities at each mutated position.

Both the native and the shuffled structures were then energy-minimized with all six force field variants (original AMBER99SB and the five reparametrized variants). For each minimized structure we computed the backbone-only RMSD to the corresponding experimental backbone. For each protein and each variant we then computed the paired difference (shuffled minus native RMSDs): a positive value indicates that the native sequence relaxes to a backbone conformation closer to the experimental structure than its shuffled counterpart.

Across all tested force field variants, the median RMSD difference remained consistently positive (Supplementary Fig. S4), demonstrating that the reparametrization preserves the ability of the force field to distinguish native sequences from randomized alternatives.

## Software and availability

All the code is available in the git repository https://github.com/grogdrinker/implicit_forcefield_reparametrization

## Acknowledgments

GO is funded by the University of Montpellier and by an ANR project (project no. ANR-23-CPJ1-0061). ST is funded by CNRS. GO thanks G. Upper for the help with the graphics of Figure 1 and for the helpful discussions.

## Methods

### Datasets

All structures used in this work were prepared with a common protocol. Heteroatom records (HETATM) were removed, because the bare AMBER99SB parameter set used here does not include water or ion parameters. The resulting PDBs were then processed with PDBFixer: missing residues and missing heavy atoms were identified and rebuilt and non-standard residues were replaced by their standard counterparts. The repaired all-atom structure was used as the reference experimental conformation in all subsequent comparisons.

In this work, we used two datasets. The primary dataset, comprising amyloid fibril structures, was used for the force field reparametrization and for the majority of our analyses. The second dataset, comprising globular protein structures, was used as an independent validation set to assess whether the reparametrization preserved the fundamental physical behavior of globular proteins.

#### Amyloid fibril dataset

Amyloid fibril structures were obtained from Amyloid Explorer [17, 18], a manually curated collection of experimentally determined amyloid structures in which every entry has been standardized to contain exactly ten identical fibril layers. Standardizing the number of layers removes the heterogeneity of the deposited structures. To keep our analysis computationally tractable, we restricted the collection to assemblies containing fewer than 5000 atoms across the ten layers. The resulting dataset comprises 121 amyloid structures, ranging from 740 to 4940 atoms.

#### Globular protein dataset

To assess whether the reparametrization preserves the general description of well-folded soluble proteins, we used an independent set of 57 small globular proteins taken from the dataset of Robustelli et al. [19]. This set was never used during training and serves exclusively as an independent control.

### Structure comparison and evaluation

#### Energy minimization

Energy minimizations were performed using the local energy minimizer implemented in OpenMM [15] (function simulation.minimizeEnergy). OpenMM supports the loading of force field parameters from XML files, allowing the minimizer to be used with arbitrary reparametrizations of the AMBER force field.

The structure was minimized to a force tolerance of 1 × 10^− 3^, with a maximum of 100 000 iterations per minimization. Because a single minimization does not always converge to a stationary point, the minimization was repeated up to five times. After each iteration, the root-mean-square (RMS) force was monitored, and the procedure was terminated when successive minimizations no longer resulted in an appreciable (< 0.1%) decrease in the RMS force. This iterative convergence criterion was used to ensure that the final structure corresponded to a stable local minimum.

#### Molecular dynamics simulations

Molecular dynamics simulations followed the same OpenMM-based procedure described above for energy minimization. OpenMM allows force field parameters to be loaded from XML files, enabling simulations with arbitrary reparametrizations of the AMBER force field. Simulations were therefore performed using the standard OpenMM simulation pipeline, with trajectory propagation carried out using simulation.step. Prior to each simulation, a single local energy minimization was performed using a force tolerance of 1.0 and a maximum of 1000 iterations. The resulting relaxed coordinates were then propagated using Langevin dynamics [20] at 300 K, with a friction coefficient of 1 ps^− 1^ and an integration timestep of 2 fs. Each trajectory consisted of 250 000 integration steps, corresponding to 0.5 ns of simulation time. Whenever possible, default OpenMM parameters were used.

#### Structure comparison

Throughout the analysis, structural agreement between simulated or energy-minimized conformations and the corresponding experimental structures was quantified using the root-mean-square deviation (RMSD) over the heavy atoms after optimal rigid-body superposition via the Kabsch algorithm [21]. This comparison is used to assess how closely the final conformation obtained from a simulation or energy minimization reproduced the experimentally determined structure. The same procedure was used to define the loss function during force field reparametrization. For each training structure, the energy-minimized conformation obtained using the current parameter set was optimally superimposed onto the corresponding experimental PDB structure, and the resulting RMSD was used as a measure of structural similarity. The loss function was then calculated from these RMSD values across the training set, quantifying the agreement between the energy-minimized structures generated by a given parameterization and their corresponding experimental structures. For the decoy analysis (“Sequence specificity: shuffled-sequence decoys”), only backbone atoms were used.

### Bilevel formulation of the force field reparametrization problem

This section formalizes the two nested optimization problems introduced in the main text and derives, step by step, the gradient that is used to train the force field parameters. We first define the problem in the specific context of force field reparametrization (the inner and outer loops), and then show how and why the implicit function theorem yields an analytic expression for the required gradient.

#### Definition of the trainable AMBER parameters

The first step is to set up a method to modify the parameters of the AMBER force field. To make the force field trainable while keeping every optimized quantity anchored to its physical meaning, the parameters are not replaced, but rescaled by multiplicative factors. Every trainable factor is a dimensionless multiplier initialized to 1.0, so that at the beginning of training the model reproduces AMBER99SB exactly. The complete set of these multipliers constitutes the trainable parameter vector ***θ***. This construction guarantees that a parameter reported as “unchanged” corresponds to a multiplier of exactly 1.0, and that the percentage deviation of any parameter from its original AMBER99SB value is directly read off from its multiplier. These multipliers are truncated so that they cannot adopt negative values.

In total, the parameter vector ***θ*** comprises 918 multipliers, of which 375 were optimized in this work. These 375 multipliers correspond to forces involved in the atom types that are present in the amyloid dataset used for training.

All parameters defined in the AMBER99SB configuration file were taken into consideration, with two exceptions: dihedral-connected parameters and 1-4 nonbonded interactions. These parameters were excluded from the optimization because modifications to them resulted in numerical instabilities during the simulations. They were therefore kept fixed at their original AMBER99SB values throughout the reparametrization. The complete list of optimized and fixed parameters can be obtained directly from the XML force field configuration file provided in the Git repository of this work.

#### Problem definition: force field reparametrization as a bilevel program

We are given the force field *F* with tunable parameters ***θ*** and a collection of experimentally determined structures. For a single structure, let **X**^exp^ ∈ ℝ^3*N*^ denote its experimental coordinates. The goal of the reparametrization is to find the parameters under which each experimentally observed structure sits at a minimum of the energy surface. This goal gives rise to two coupled optimization problems, which we call the inner and outer problems.

##### Inner problem (energy minimization)

For a *fixed* parameter set ***θ***, the force field defines a relaxed structure through energy minimization. Starting from a set of coordinates, the relaxed conformation is the minimizer of the potential energy.

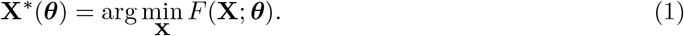

The relaxed structure is written **X**^∗^(***θ***) to emphasize that it is a *function of the parameters*: changing ***θ*** changes the energy surface and therefore moves the location of its minimum. This is precisely the standard energy minimization performed by a molecular-mechanics engine such as OpenMM. Being a minimum, **X**^∗^(***θ***) satisfies the first-order optimality (stationarity) condition that the energy gradient with respect to the coordinates is equal to zero. This means that the forces acting on every atom (minus the gradient of the energy with respect to the atom positions) are (approximately) zero:

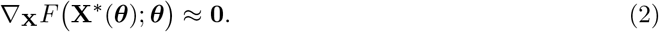

The inner problem (1) was solved by energy minimization with OpenMM, using the current parameter set ***θ***. This stationarity condition is required to compute the gradient, as it allows us to treat the inner minimization as a black box that returns a relaxed structure for a given ***θ***. This enables the use of OpenMM for energy minimization without differentiating through each minimization step.

##### Outer problem (parameter fitting)

The parameters ***θ*** are then adjusted so that the relaxed structure **X**^∗^(***θ***) is as close as possible to the experimental one **X**^exp^. Closeness is measured by the RMSD between the two sets of coordinates, after the application of the Kabsch algorithm. This value serves as the outer objective:

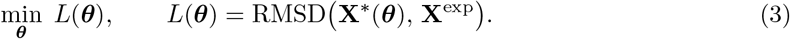

#### Gradient of the outer objective via the implicit function theorem

To optimize the parameters by gradient descent we need the gradient of the outer objective, ∇_***θ***_*L*. The thing that makes the problem difficult is the fact that the two problems are nested: evaluating the outer objective *L*(***θ***) requires first solving the inner problem (1) to obtain **X**^∗^(***θ***). The essential difficulty is that the outer variable ***θ*** enters *L* only *indirectly*, through the minimizer **X**^∗^(***θ***), which is itself defined as the outcome of an optimization and is not available in closed form.

In the next paragraphs we show the steps to apply the implicit function theorem to remove this obstacle and turn the issue into a linear-algebra problem involving only quantities evaluated at the converged structure.

##### Step 1: chain rule through the relaxed structure

Recall that the outer objective (Eq. (3)) depends on the parameters only through the relaxed structure **X**^∗^(***θ***).

Because ***θ*** reaches *L* only through the intermediate coordinates **X**^∗^(***θ***), a perturbation of a single parameter *θ*_*j*_ shifts every coordinate at once, and the loss responds to all of these shifts together. The multivariable chain rule collects these contributions as a sum over the 3*N* coordinates:

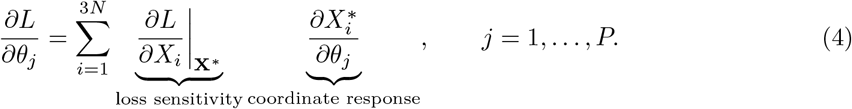

The first factor, *∂L/∂X*_*i*_ evaluated at **X**^∗^, is how the RMSD changes when coordinate *i* is displaced. The second, 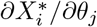, is how coordinate *i* of the minimizer moves when parameter *j* changes.

Collecting these scalar objects into vectors and matrices, we define the coordinate gradient of the loss and the Jacobian of the minimizer,

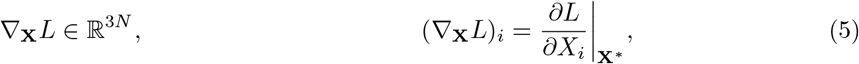

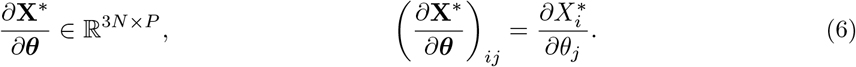

The sum in (4) is then a single matrix–vector product. Since the output index *j* of ∇_***θ***_*L* ∈ℝ^*P*^ labels *parameters* while the summation index *i* labels *coordinates*, the Jacobian enters transposed, mapping the coordinate-space gradient ∇_**X**_*L* ∈ ℝ^3*N*^ into the parameter-space gradient ∇_***θ***_*L* ∈ ℝ^*P*^ :

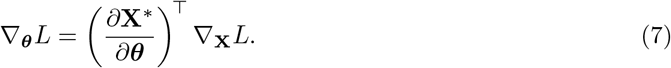

The dimensions confirm the transpose: (*P*×3*N*) · (3*N*) = (*P*). Equation (7) splits the required gradient into an explicit, easily evaluated factor ∇_**X**_*L*, immediate because the RMSD is a known closed-form function of the coordinates, and an implicit factor *∂***X**^∗^*/∂****θ***, the Jacobian of the minimizer, which has no closed form and requires the application of the implicit function theorem.

##### Step 2: the minimizer is defined implicitly by the stationarity condition

Although **X**^∗^(***θ***) has no closed form and is only available as the output of a numerical minimization, it satisfies a known condition at every ***θ***: the stationarity condition (2). We therefore obtain the Jacobian from this condition. Define the residual map

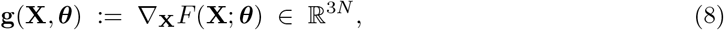

so that the stationarity condition reads **g X**^∗^(***θ***), ***θ*** = **0** for all ***θ***. This is exactly the setting of the implicit function theorem: **g** is continuously differentiable and, provided its Jacobian with respect to **X** is non-singular at the solution, the equation **g**(**X, *θ***) = **0** defines **X**^∗^(***θ***) as a differentiable function of ***θ*** in a neighbourhood of that solution.

##### Step 3: differentiate the stationarity identity

The strategy is to differentiate the identity **g X**^∗^(***θ***), ***θ*** = **0** itself, rather than **X**^∗^(***θ***) directly. Because the identity holds for *every* ***θ***, its total derivative with respect to ***θ*** must also vanish. The map **g** depends on ***θ*** in two ways: directly through its second argument, and indirectly through **X**^∗^(***θ***). Applying the chain rule to both channels gives

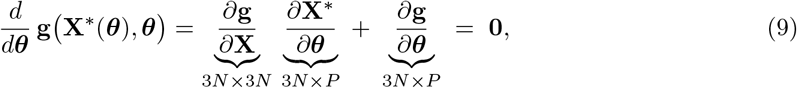

where the first term is the indirect contribution (through the minimizer, which itself moves with ***θ***) and the second is the direct one (the explicit parameter dependence).

Because **g** = ∇_**X**_*F*, both derivatives of **g** are second derivatives of the energy:

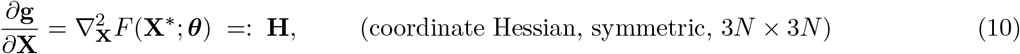

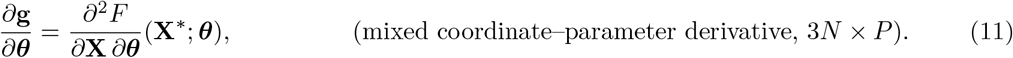

Both terms have a direct physical meaning. The Hessian **H** is the curvature of the energy at the minimum, i.e. how the force on each atom changes when the atoms themselves are displaced. The mixed derivative *∂*^2^*F/∂***X** *∂****θ*** measures how the force on each atom changes when a force field parameter is perturbed, and it is the only channel through which the parameters act on the relaxed structure.

##### Step 4: solve for the Jacobian

The implicit function theorem requires that **H** is invertible at the minimizer, so Eq. (9) can be solved for the unknown Jacobian:

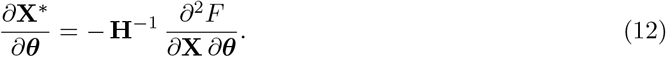

##### Step 5: assemble the outer gradient

Substituting the Jacobian (12) into the chain rule (7) and using the symmetry of **H** (**H**^⊤^ = **H**, hence **H**^− ⊤^ = **H**^− 1^) gives the gradient of the outer objective in closed form,

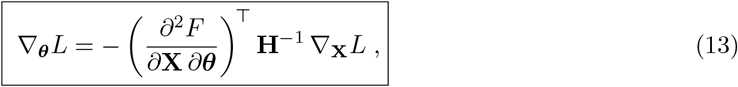

with 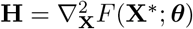 and all quantities evaluated at the relaxed structure **X**^∗^.

Two properties of Eq. (13) are decisive in practice. First, every term is evaluated only at the converged structure **X**^∗^: the coordinate gradient of the RMSD ∇_**X**_*L*, the coordinate Hessian of the energy **H**, and the mixed second derivative, *∂*^2^*F/∂***X** *∂****θ***. The expression makes no reference to the path the minimizer followed to reach **X**^∗^. The outer gradient therefore depends only on the returned stationary point, not on the trajectory (or algorithm) used to reach it, and the inner minimizer can be treated as a black box that merely returns a relaxed structure for a given ***θ***, which is what allows a highly optimized external engine (OpenMM) to be used for the inner problem without differentiating through it. Second, the derivation relies on the stationarity condition in Eq. (2): Equation (13) is valid only if **X**^∗^ is sufficiently close to a stationary point, such that ∇_**X**_*F* (**X**^∗^; ***θ***) ≈ **0**.

#### Matrix-free evaluation of the gradient

Equation (13) was not evaluated by forming or inverting the Hessian **H**, a 3*N* × 3*N* matrix that is intractable at the system sizes considered here. Instead, it was evaluated in a matrix-free fashion, following the standard adjoint approach to implicit differentiation [13]. The energy *F*, its coordinate gradient∇_**X**_*F*, and all higher derivatives were implemented in PyTorch, so that every required quantity is obtained by automatic differentiation.

Rather than building the full Jacobian *∂***X**^∗^*/∂****θ***, which would require one linear solve with **H** for each of the *P* parameters, we first solve a single linear system for the adjoint vector **v**,

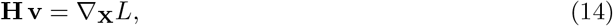

and then obtain the parameter gradient in one additional automatic-differentiation pass,

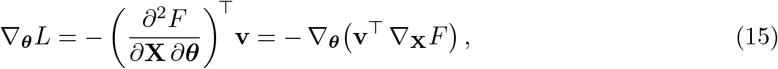

where **v** is held constant. The cost of the gradient therefore amounts to one linear solve, independently of the number of parameters.

The system (14) was solved with the conjugate-gradient (CG) method, which accesses **H** only through Hessian–vector products. These were computed exactly by automatic differentiation, without ever building **H** [22]. Following common practice for implicit gradients [23], the system was damped, (**H** + *µ***I**) **v** = ∇_**X**_*L* with *µ* = 1.0. The solver used up to 200 iterations and a residual tolerance of 1 × 10^− 8^ If the relative residual exceeded 0.01, the solve was repeated with *µ* = 10.0.

##### Outer-loop optimization and training schedule

For each structure the inner minimization (1) was solved with the current ***θ***, the outer RMSD (3) was evaluated, its gradient with respect to ***θ*** was accumulated through Eq. (13), and the optimizer step was applied. The parameters ***θ*** were updated with each structure (batch = 1) with the Adam optimizer [24] at a learning rate of 0.001. Before each update, the gradient norm was clipped to a maximum of 1.0. One training epoch consisted of a full pass over all 121 amyloid structures. Training was run for 10 epochs. The trained multipliers were exported to an OpenMM-compatible XML parameter file, so that the reparametrized force field can be used directly in standard OpenMM energy minimizations and molecular-dynamics runs.

##### Regularization

To discourage unnecessarily large departures from the original physical parameterization, we added an L2 regularization term (“weight decay” [25]) that penalizes the deviation of each multiplier from its initial value of 1.0. Denoting the multipliers collectively by ***θ***, the regularized objective per structure is

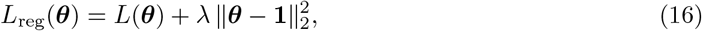

so that small parameter changes are only weakly penalized while increasingly large deviations are progressively discouraged. In practice the penalty was applied at the gradient level, adding 2*λ*(***θ***^−^**1**) to the outer gradient before each optimizer step. The regularization strength *λ* was swept over the values *λ* ∈ {0.005, 0.01, 0.05, 0.1}, together with the unregularized case *λ* = 0. All other training settings were held fixed across the sweep.

##### Cross-validation procedure

To test whether the reparametrization generalizes to structures not seen during training, we performed a grouped five-fold cross-validation. To prevent near-duplicate structures (sequence polymorphs, or different resolved ranges of the same protein) from biasing the training, the folds were defined by grouping rather than by a random split. For each structure, the longest distinct chain sequence was taken as the representative monomer sequence. Structures were then clustered by substring containment: two structures were assigned to the same cluster whenever one representative sequence is a contiguous substring of the other. This places every truncation variant and resolved range of a given protein in a single cluster, reducing the 121 structures to 39 clusters. It is important to note that amyloid fibrils formed by the same sequence can adopt markedly different structures, a phenomenon known as structural polymorphism [26]. Whole clusters were distributed over the five folds by a size-balancing greedy rule (largest cluster first, assigned to the currently smallest fold), so that clusters were never split and no protein, nor any other member of its family, appeared in both the training and the held-out set. This produced folds of 40, 21, 20, 20 and 20 structures.

For each fold, the force field was retrained from scratch on the other four folds and evaluated on the test fold, using the same procedure and hyper-parameters as the full-dataset training and without regularization (*λ* = 0).

#### Software and implementation

The molecular-mechanics force field, its differentiable (multiplier-based) parameterization, the analytic energy and gradient, the Hessian-vector products and the implicit-function-theorem gradient were implemented in PyTorch. Energy minimizations and molecular-dynamics simulations of the reference and reparametrized force fields were performed with OpenMM. Figure components have been done using biorender [27], pymol [28] and chimeraX [29].

## Notes

https://github.com/grogdrinker/implicit_forcefield_reparametrization/

### Competing Interest Statement

The authors have declared no competing interest.

