## Supplementary Figures for "The (implicit) force is with you: an application of the implicit function theorem for automatic protein force fields reparametrization"

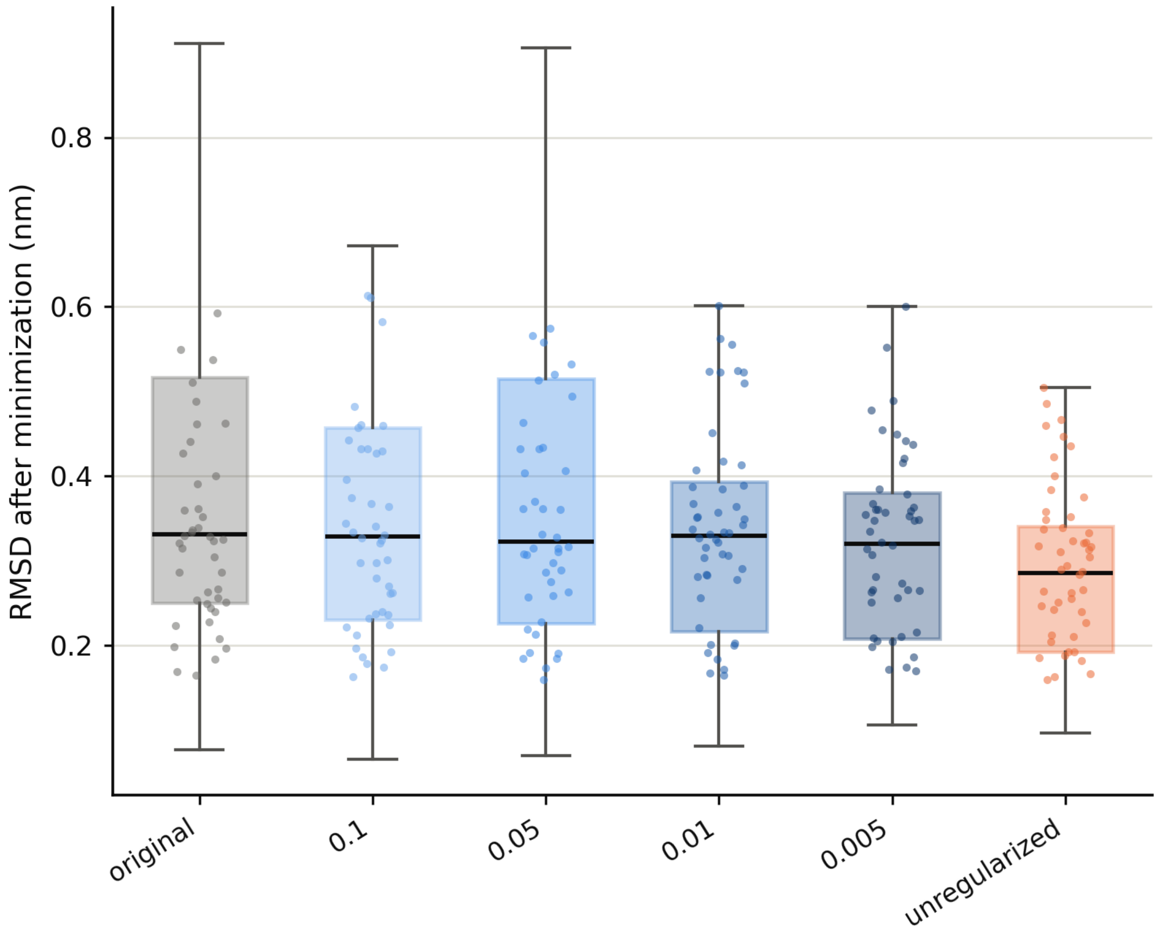


**Figure S1:** Energy minimization of non-amyloid proteins. The boxplots show the RMSD distribution obtained after energy minimization of a dataset of 57 globular proteins using the original AMBER99SB force field and the five reparametrized variants. RMSD values were calculated using Kabsch alignment over the heavy atoms between the minimized conformations and the corresponding experimentally determined structures. The RMSD distributions are highly similar across all force field variants, including the original AMBER99SB parameterization, indicating that the reparametrization optimized on amyloid structures does not compromise the general performance of the force field on unrelated globular proteins. Scatter points are displayed after clipping to the 10th–90th percentile range for visualization purposes only. All statistical analyses were performed on the complete dataset.

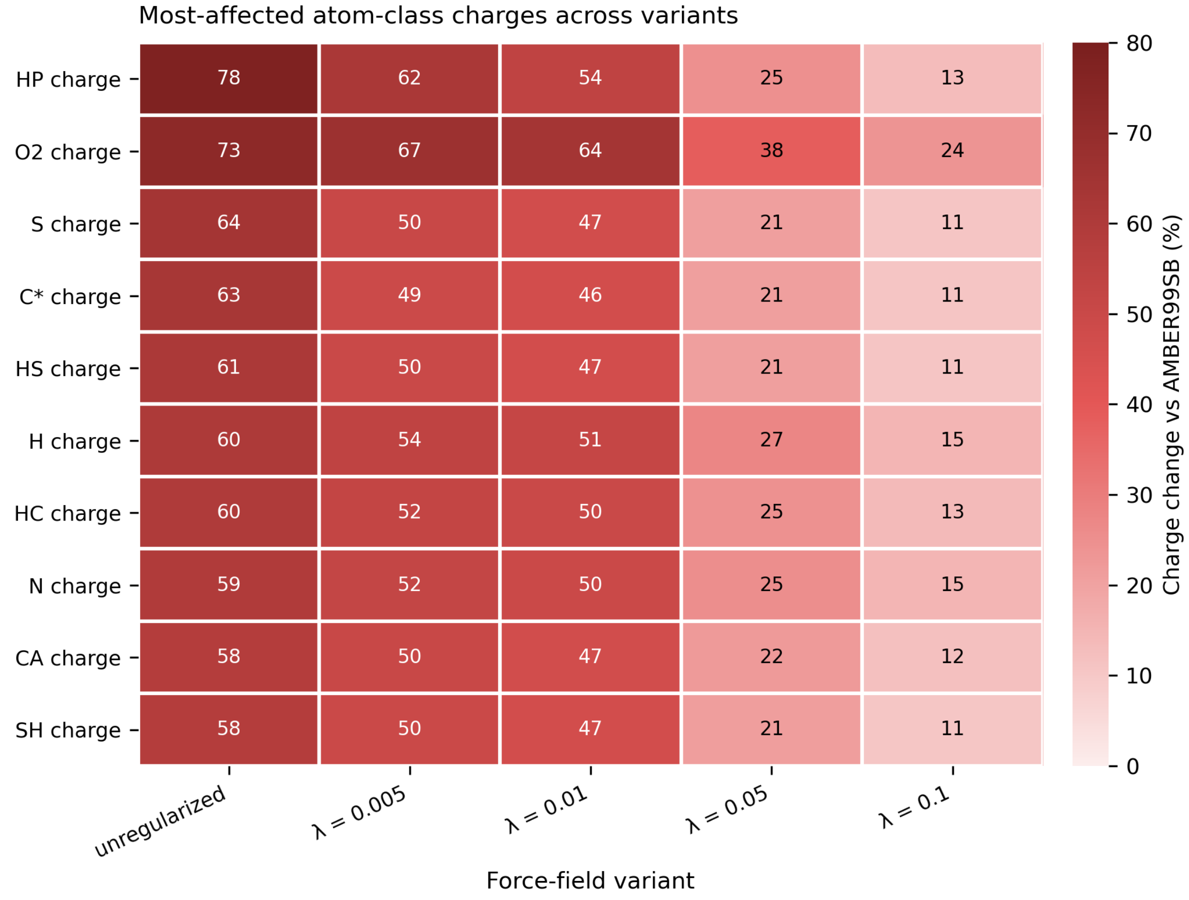


**Figure S2:** Top force-field parameter changes across variants. Per-atom-class charge change (% relative to the original AMBER99SB value) for the ten most-affected atom classes, shown for the unregularized model and each regularization strength.

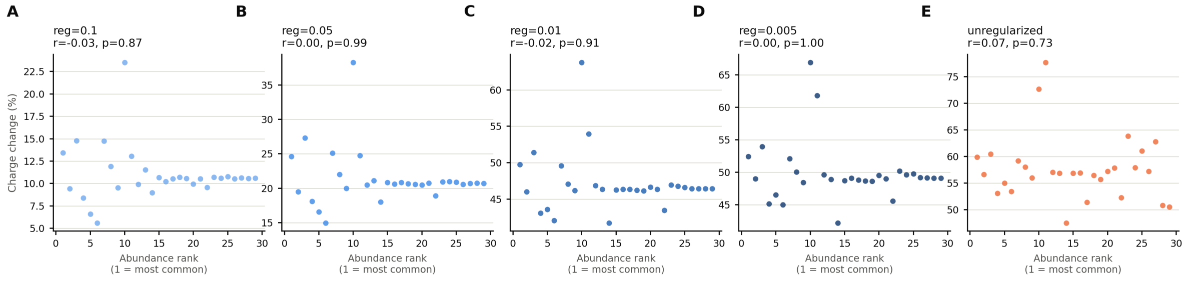


**Figure S3:** Correlation between atom class abundance and charge variation during reparametrization. The plots show the relationship between the abundance of each atom class in the amyloid dataset and the magnitude of the corresponding charge change after reparametrization. Each panel corresponds to a different regularization strength ($\lambda$). No significant correlations are observed for any of the tested conditions, with correlation coefficients consistently close to zero across all variants. These results indicate that atom class abundance is not a confounding factor in the IFT-based reparametrization procedure and that charge updates are not driven by the frequency of occurrence of specific atom types.

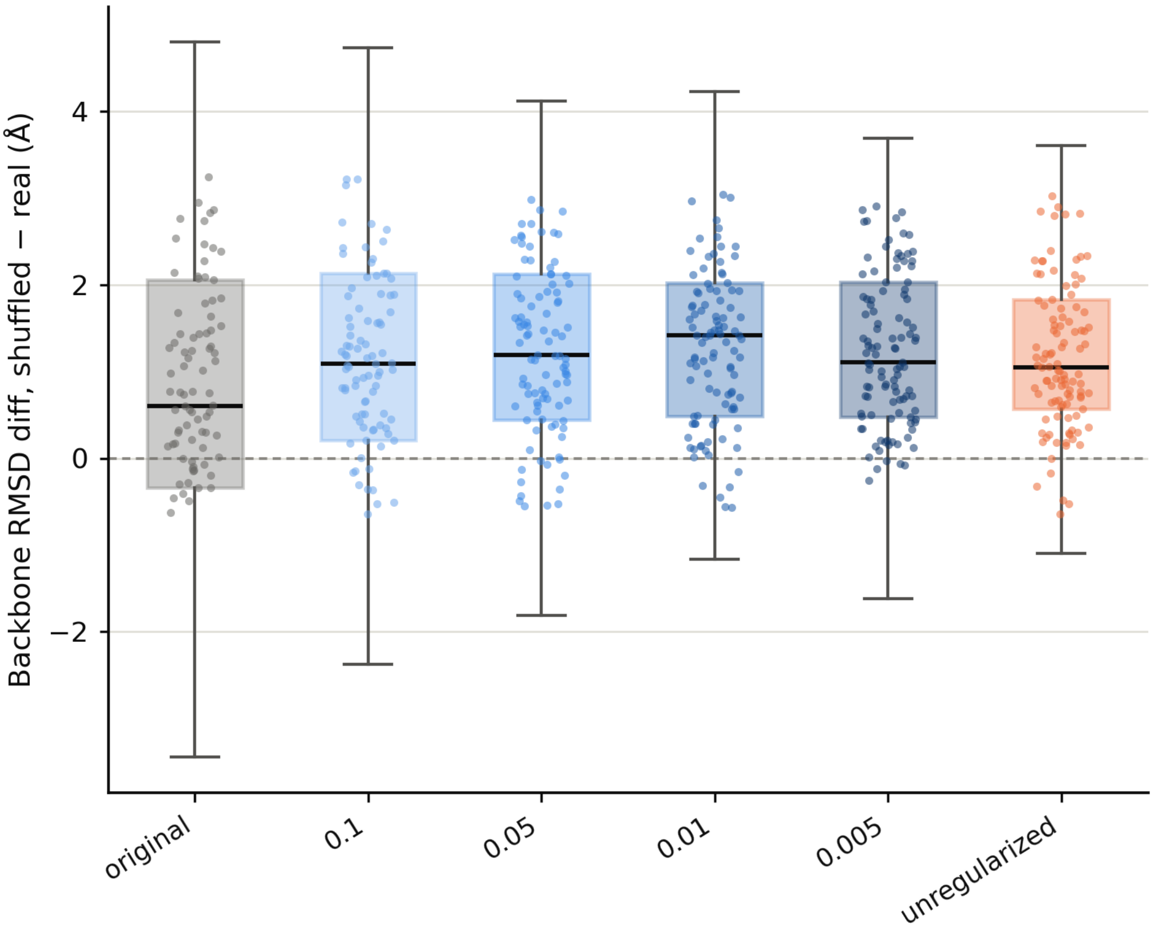


**Figure S4:** Sequence specificity of force field reparametrization. For each of 121 amyloid structures, a shuffled-sequence decoy was generated and independently energy-minimized under all 6 force-field variants. Each point/box shows, per protein, the difference in backbone-only (N, C$\alpha$, C, O) Kabsch RMSD to the original experimental structure between the shuffled decoy and the real sequence (shuffled − real). Values above zero indicate the real sequence relaxes closer to the native backbone conformation than its shuffled counterpart. All 6 variants show a positive median (0.60 Å for the original AMBER force field and approximately 1.0–1.4 Å for the reparametrized variants), indicating every tested force field discriminates the native sequence from its shuffled decoy, with the reparametrized variants discriminating at least as strongly as the original. Scatter points are clipped to the 10th–90th percentile for display only (statistics are computed on the full data).
